# Nitrous Oxide Enhances Focused Ultrasound Mediated Delivery of Viral Gene Therapy

**DOI:** 10.1101/2021.07.11.451978

**Authors:** Bhavya R. Shah, Rachel M. Bailey, Ibrahim Youssef, Sydni K. Holmes, Austin Marckx, Sandi Jo Estill-Terpac, Marc Diamond, Rajiv Chopra

## Abstract

Transcranial Magnetic Resonance Guided Focused Ultrasound (MRgFUS) can be used in conjunction with intravenous microbubbles to open the blood brain barrier (BBB) in discrete brain regions. This method is limited by the microbubble dose, the amount of energy that can be safely delivered across an intact skull, and the inability to target areas of the brain outside of central brain regions. We find that N_2_O administration substantially decreases the required dose of microbubbles and/or focused ultrasound pressure while still reliably opening the blood brain barrier. Additionally, we observe that the use of nitrous oxide improves the delivery of adeno-associated virus (AAV9) vector when compared to medical air. Our findings may help overcome focused ultrasound limitations and improve targeted delivery of therapeutics in humans.

**One Sentence Summary:** Nitrous Oxide potentiates Blood Brain Barrier opening with Focused Ultrasound and enhances viral vector delivery.

## Introduction

The blood brain barrier (BBB) hinders the delivery of diagnostic and therapeutic agents to the brain. Attempts to bypass the BBB have included intra-arterial hyperosmotic solutions,(*1, 2*) targeting cell surface receptors, (*3, 4*) and intraventricular or intrathecal delivery. However, none of these methods target discrete brain regions. Although stereotactic injection can overcome these limitations, it is limited by a risk of hemorrhage, injury to surrounding cells, and limited spread of the therapy from the site of delivery.

Transcranial Magnetic Resonance Guided Focused Ultrasound (MRgFUS) is a novel, non-invasive, image-guided procedure that opens the blood brain barrier (BBB) in discrete brain regions. It is a unique approach to non-invasively deliver therapies to discrete brain regions. After an intravenous infusion of microbubbles, ultrasound waves are focused across an intact skull onto a specific brain region. In this targeted region, the intravascular microbubbles oscillate and disassemble tight junctions, which results in temporary opening of the BBB. (*5-7*) The safety and feasibility of MRgFUS BBB opening is being established in patients with neurodegenerative diseases and brain tumors. (*8-10*) Equally exciting is the opportunity to deliver gene therapy including Designer Receptors Exclusively Activated by Designer Drugs (DREADDs) across the BBB. (*11*) While initial clinical trial data is promising, optimization of parameters to account for variability in skull thickness, regional vascularity, vessel diameter, and cerebral blood flow are critical elements in successful human translation. Improving the reproducibility of BBB opening with the lowest microbubble dose and FUS energy possible is a desirable goal prior to widespread clinical adoption.

Nitrous Oxide (N_2_O) is a medical gas that is widely used in the outpatient setting and produces mild analgesia (*5*). We find that N_2_O reduces the FUS energy and microbubble dose required to open the BBB. T1-weighted postcontrast MR imaging (gadolinium) is commonly used to target and confirm BBB opening with FUS (*12, 13*). However, post-contrast MRI does not provide real-time feedback during sonication. Real-time monitoring of acoustic emissions from stimulated microbubbles can serve as a feedback control mechanism during FUS treatment (*8, 14-18*). Real-time feedback during sonication enables the user to determine if sufficient energy is being delivered to open the BBB or if too much energy is causing injury. Safe, stable microbubble oscillations emit harmonic emissions that consist of subharmonic, ultra-harmonic, and second harmonic emissions (*19*). However, as the acoustic pressure is increased and collapsing microbubbles undergo cavitation, they emit broadband acoustic emissions (*16*). Monitoring of real-time harmonic emissions makes it possible to overcome variability that arises from differences in skull thickness, regional vascularity, vessel diameter, and cerebral blood flow. Differences in vascularity and blood flow can produce substantial spatial and temporal variations in local concentration of microbubbles which leads to inconsistency when determining the optimal amount of focused ultrasound energy needed to open the BBB.(*16, 20-24*) Optimizing and monitoring interactions between microbubbles and FUS is essential to safe and reproducible BBB opening in humans.

We find that when administering N_2_O, microbubble harmonic emissions are more reliably detected, there is more consistent and reproducible BBB opening on postcontrast T1-weighted MRI, and we can either reduce microbubble doses or FUS pressures to achieve BBB opening. We also observe that this method improves the delivery of an adeno-associated virus (AAV) vector to discrete brain regions.

## Results

### Nitrous Oxide Amplifies Subharmonic Acoustic Emissions

Since safe BBB opening can be monitored with real-time microbubble acoustic monitoring, we first sought to study the effect of N_2_O on stimulated microbubble acoustic emissions. When N_2_O was present in the blood, a reliable, sharp subharmonic emission could be detected (Fig. 1A). However, this subharmonic emission was substantially less conspicuous when medical air was used (Fig. 1A). To quantitatively compare the effect of N_2_O on the subharmonic emission, an area under the curve (AUC) was defined as the sum of the harmonic amplitudes in the frequency spectra centered at the subharmonic of the transmit frequency for all bursts at a single pressure and in a single exposure location. At all pressures we find a statistically significant increase in the mean AUC when N_2_O was used instead of medical air (Fig. 1B). The amplification factor of the mean AUC with N_2_O when compared to medical air at a FUS pressure of 0.392MPa is 50.

**Fig. 1.**
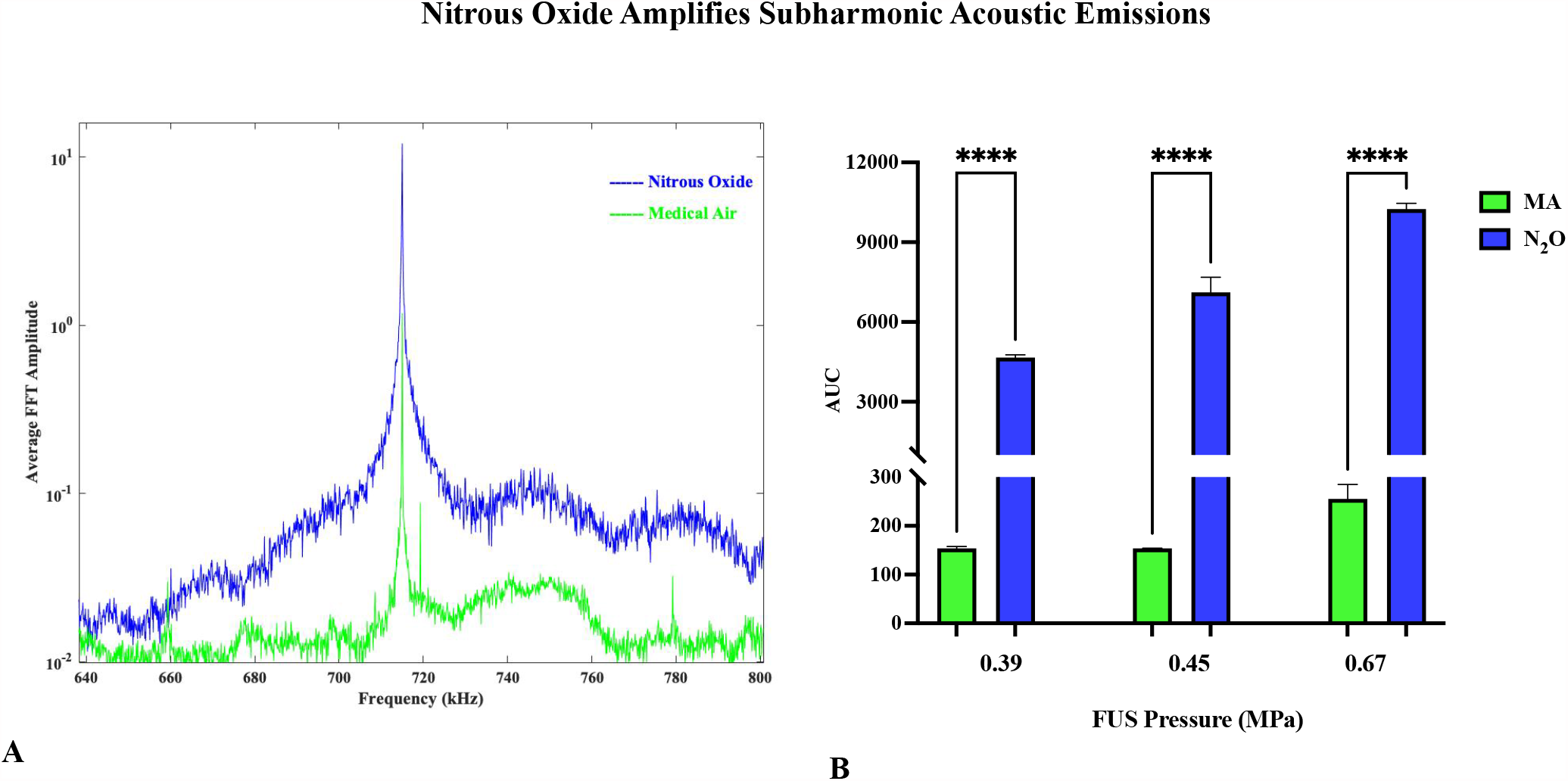
Nitrous Oxide Amplifies Subharmonic Acoustic Emissions. (A) Comparison of Average AUC Spectra with Nitrous Oxide and Medical Air (B). There is a statistically significant increase in the mean AUC when N_2_O was used instead of medical air

### Nitrous Oxide Enhances BBB opening as determined on post-gadolinium T1 weighted images

Since BBB opening can be confirmed with post-gadolinium T1 weighted images (T1WI), we then sought to understand the effect of N_2_O on enhancement in mice. When FUS pressures, microbubble concentrations, and gadolinium doses are kept constant, there is substantially more enhancement on post-contrast T1WI when BBB opening was performed with N_2_O instead of medical air (Fig. 2). To quantitatively differentiate the effect of N_2_O vs medical air on enhancement, we calculated Enhancement Rate (Equation 1) based on the T1-weighted MR images.

**Fig. 2.**
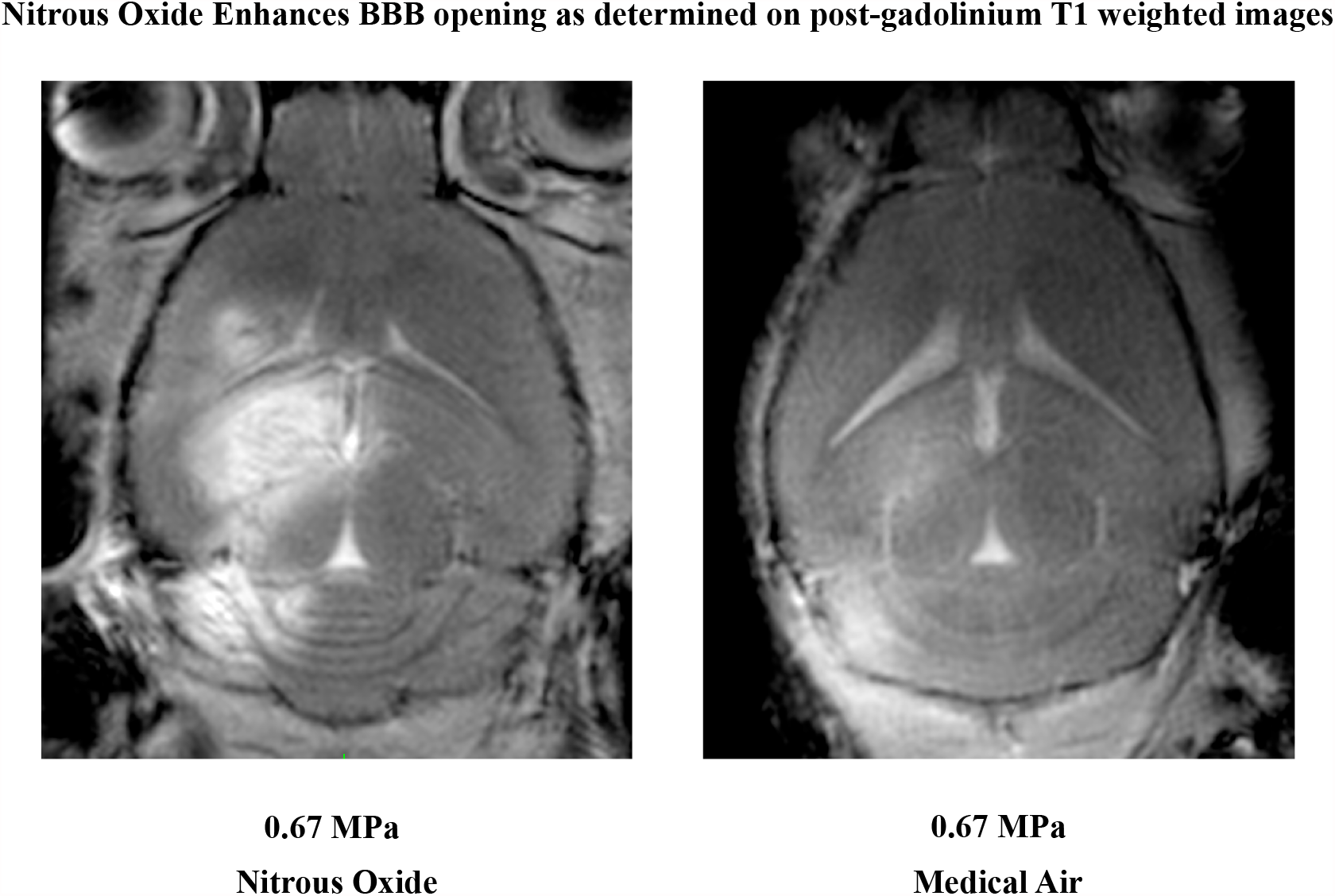
Nitrous Oxide Enhances BBB opening as determined on post-gadolinium T1 weighted images. T1W MR image demonstrating Gadolinium leakage after BBB opening with at 0.67 MPa A. Medical Air and B. Nitrous Oxide at 0.67 MPa. There is more qualitative contrast enhancement after BBB opening in the striatum, hippocampus, and cerebellum with Nitrous Oxide than with Medical Air.

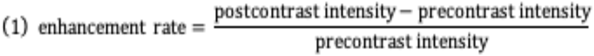

There is a statistically significant improvement in enhancement rate after BBB opening with N_2_O verses medical air (Fig. 3). Gadolinium is too large to cross an intact BBB. Therefore, enhancement on T1 weighted images represent the degree of gadolinium deposition after BBB opening. A higher enhancement rate is indicative of more contrast deposition and improved BBB opening.

**Fig. 3.**
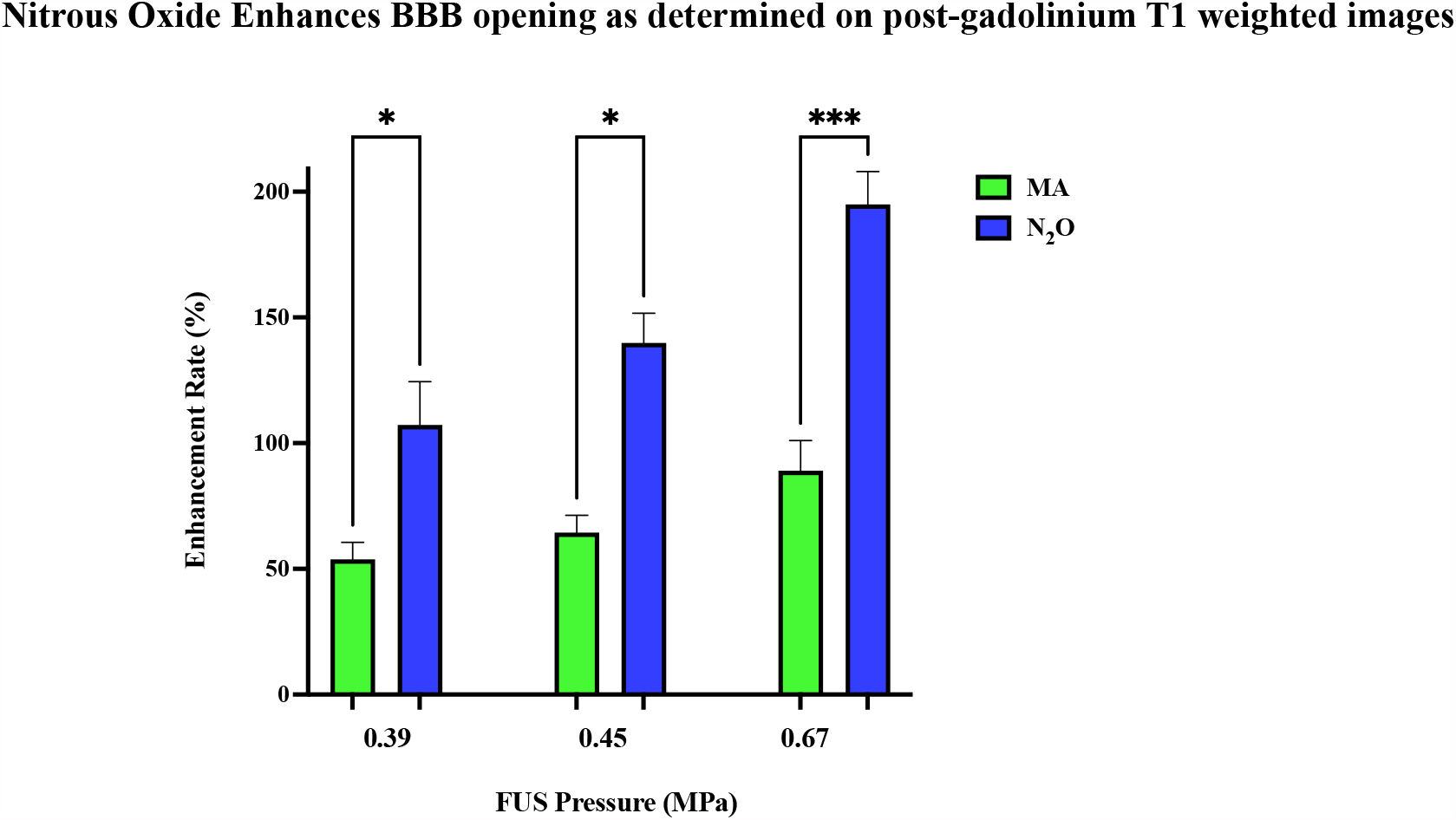
Nitrous Oxide Enhances BBB opening as determined on post-gadolinium T1 weighted images. There is a statistically significant increase in Enhancement Rate after BBB opening with nitrous oxide than with medical air at FUS pressures ranging from 0.39 MPa-0.67 MPa.

### Nitrous Oxide reduces FUS pressures required to open the Blood Brain Barrier

Previous studies by other groups have defined 0.30 MPa as the lower limit of FUS pressures required for in-vivo BBB opening.(*25, 26*) However, when N_2_O is administered, reliable AUCs can be detected at 0.280 MPa (Fig. 4A). Additionally, the Enhancement Rate is comparable to medical air at a much higher pressure (Fig. 4B).

**Fig. 4.**
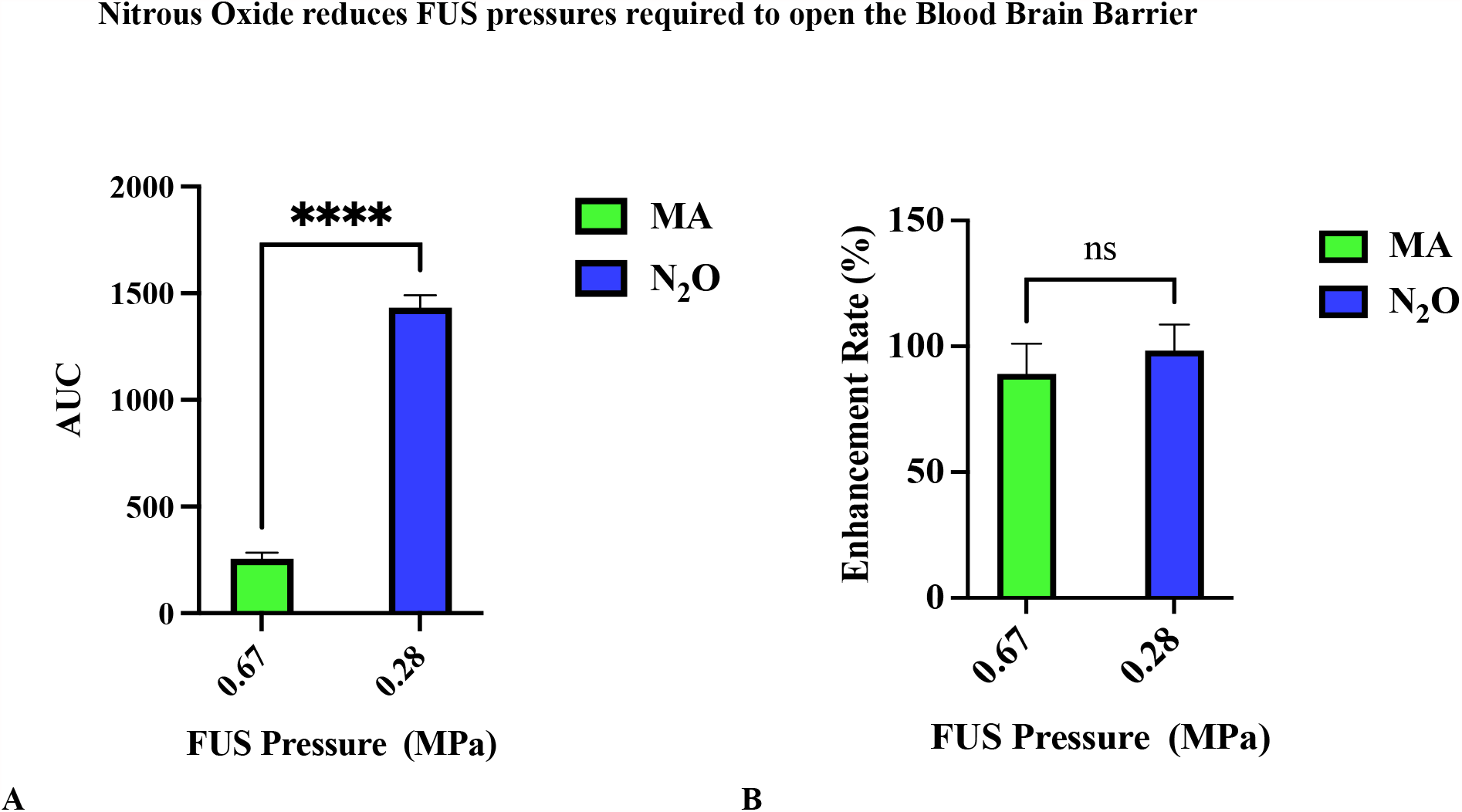
Nitrous Oxide reduces FUS pressures required to open the Blood Brain Barrier. When using nitrous oxide and a FUS pressure of 0.28 MPa both the average AUCs (A)) and Enhancement Rates (B) are higher than with Medical Air at a pressure of 0.67 MPa. Additionally, the Enhancement Rate is comparable to medical air at a much higher pressure.

### Nitrous Oxide allows for a reduction in microbubble dose for opening the Blood Brain Barrier

Although several studies are evaluating the safety and efficacy of BBB opening in humans, a key limitation is microbubble dose constraints. When the microbubble dose is diluted to one-thousandth the clinical dose, N_2_O amplifies the AUC which is comparable to an AUC seen with medical air at much higher pressures (Fig 5A). We also find that N_2_O enables a substantial reduction in the microbubble dose required to obtain BBB opening at dilutions of a tenth to one-hundredth of the clinical dose (Fig. 5B) However, when there are not enough microbubbles in circulation (one-thousandth of the clinical dose) there is very weak enhancement and as expected poor BBB opening. As the FUS pressure is increased, the mean AUC (Fig. 6A) and (Fig. 6B) Enhancement Rate increase, even when the microbubble dose is diluted by x1000.

**Fig. 5.**
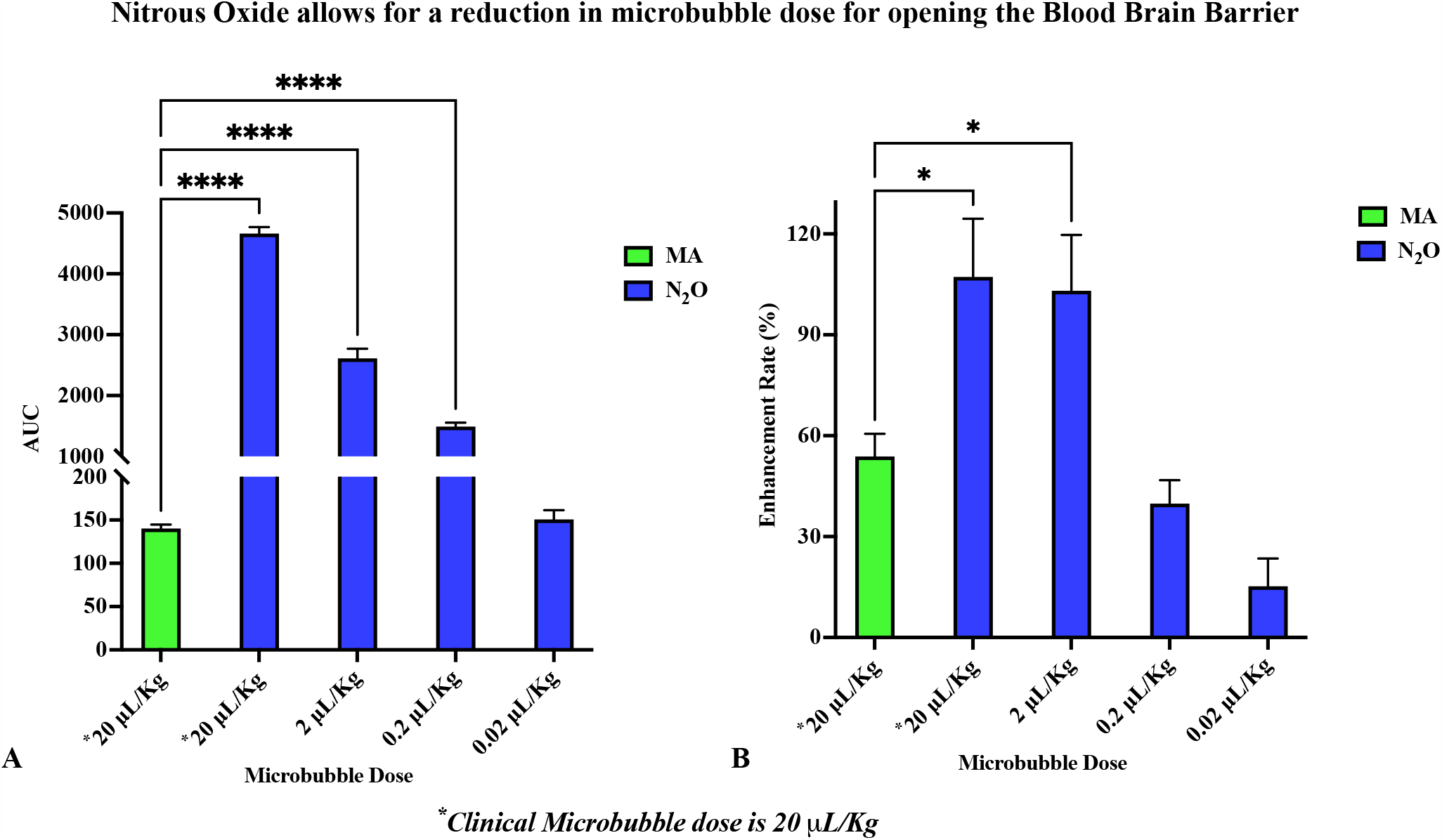
Nitrous Oxide allows for a reduction in microbubble dose for opening the Blood Brain Barrier. Effect of microbubble dilution on the mean AUC (A) and Enhancement Rate (B) at a FUS pressure of 0.39 with Medical Air verses Nitrous Oxide. When using Nitrous Oxide, the microbubble dose can be diluted 1000x and still give a mean AUC similar to Medical Air with a clinical microbubble dose. When microbubble dose is diluted 100x fold an Enhancement Rate comparable to medical air at a clinical microbubble dose is identified.

**Fig. 6.**
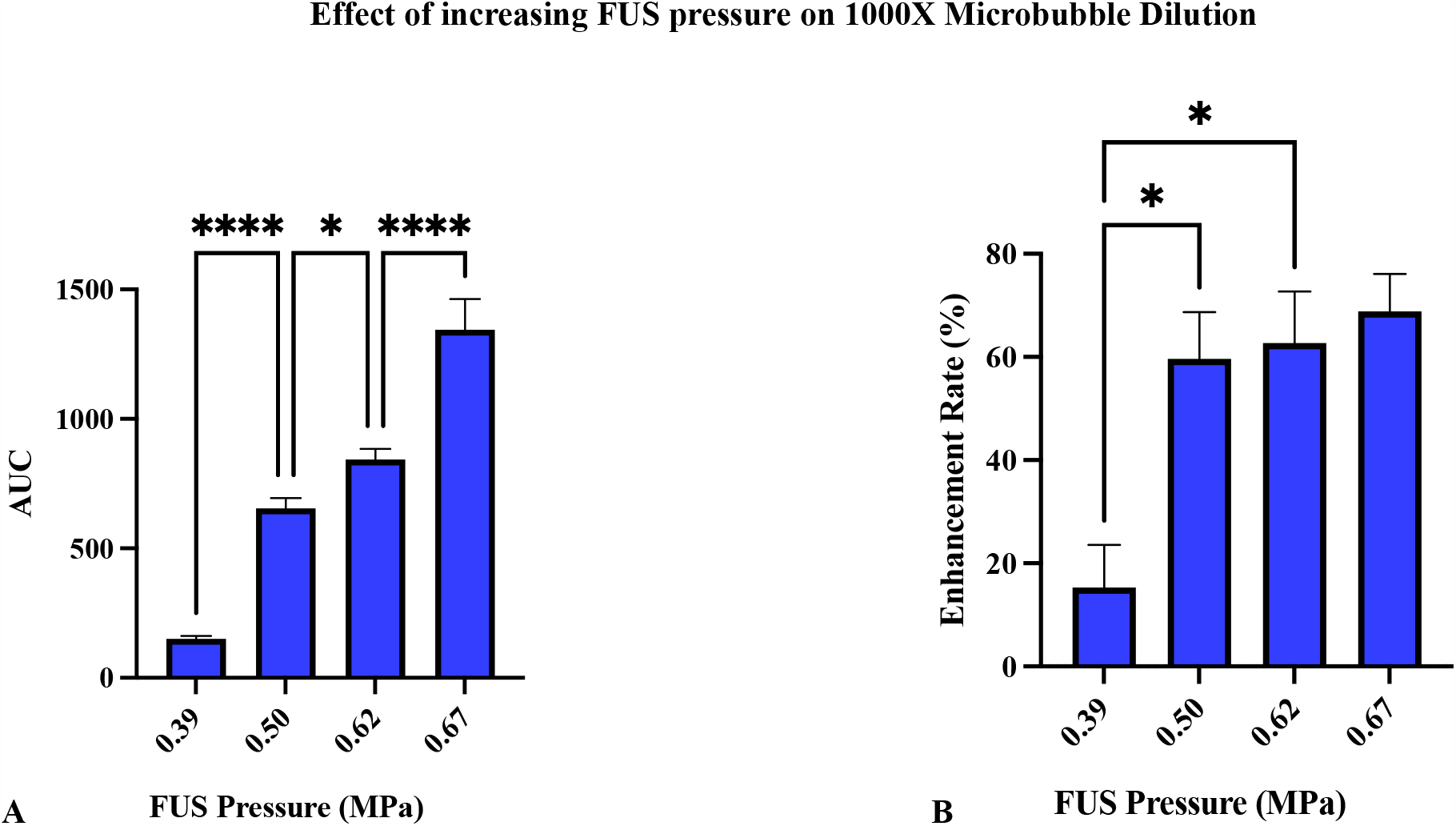
Effect of increasing FUS pressure on 1000X Microbubble Dilution. Increasing FUS pressures with 1000X dilution using Nitrous Oxide increases mean AUC (A) and Enhancement Rate (B) at even a thousand-fold microbubble dilution.

### Nitrous Oxide improves the delivery of viral gene therapy to neurons

After we determined the effects of N_2_O on microbubble stimulated acoustic emissions and enhancement rate, we sought to evaluate whether this translated into a clinically relevant bio-effect. AAV9 is one of the most frequently used vectors to deliver gene therapies to the brain due to its ability to cross the BBB and dose-dependent transduction of neural tissues. To test whether N_2_O could potentiate delivery of therapies to targeted regions of the brain, we delivered a low-dose (1E+9 vg/g) of AAV9 vector packaging a GFP transgene to the right hippocampus in mice after BBB opening with ultrasound in the presence of either medical air or N_2_O. The left hemisphere was untargeted and used to assess AAV9 transduction in the absence of FUS. Compared to medical air, we find that N_2_O allows for significantly more delivery of AAV9/GFP as demonstrated by the enhanced expression of GFP in both hippocampal neurons and astrocytes. Additionally, even at a higher pressure, 0.672 MPa, AAV9/GFP delivery with medical air was significantly less than with N_2_O with a FUS pressure of 0.392 MPa (Fig. 7A-D).

**Fig. 7.**
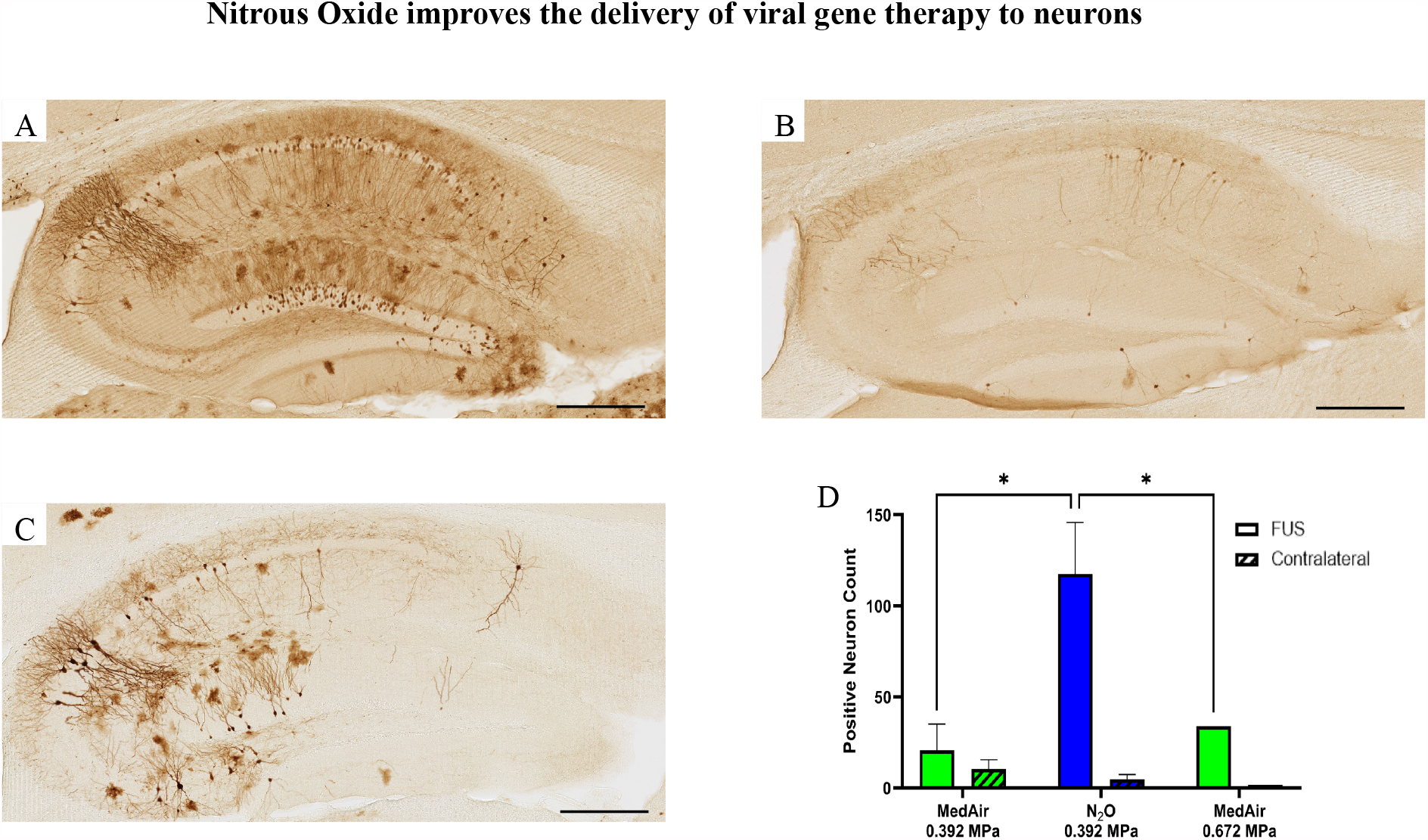
Nitrous Oxide improves the delivery of viral gene therapy to neuron. Nitrous oxide enhances gene expression following FUS-mediated delivery of scAAV9/GFP to the hippocampus. A-C Representative images showing GFP transgene staining in the hippocampus of the FUS-targeted brain hemisphere of mice following intravenous injection of 1×10^9^ vg/g scAAV9/GFP. (A) At FUS pressures of 0.392 MPa, and the use of nitrous oxide shows abundant GFP expression (B) At FUS pressures of 0.392 MPa, and the use of medical air shows limited transgene expression, while (C) Increasing the FUS pressure to 0.672 MPa while using medical air increases GFP expression as compared to a FUS pressure of 0.392 MPa medical air (B) but is still significantly less than use of nitrous oxide and 0.392 MPa (A). (D) Quantification of GFP positive neurons in the hippocampus of the FUS-treated brain hemisphere and in the contralateral, untreated brain hemisphere in each group. Data is represented as the mean ±SD. *P<0.05 [two-way ANOVA (FUS condition x brain side): main effect of FUS condition indicated]. Scale bar, 300 um.

### Opening the BBB with Nitrous Oxide at lower pressures results in less acute injury than when obtaining BBB opening using medical air at higher FUS pressures

Reliable BBB opening with medical air routinely requires FUS pressures of 0.672 MPa. When N_2_O is used, reliable BBB opening occurs between 0.280-0.392 MPa. At these lower FUS pressures, there is less microgliosis as shown by decreased Iba1 staining immediately after FUS suggesting that N_2_O mediated BBB opening is safer at lower pressures (Fig. 8).

**Fig. 8.**
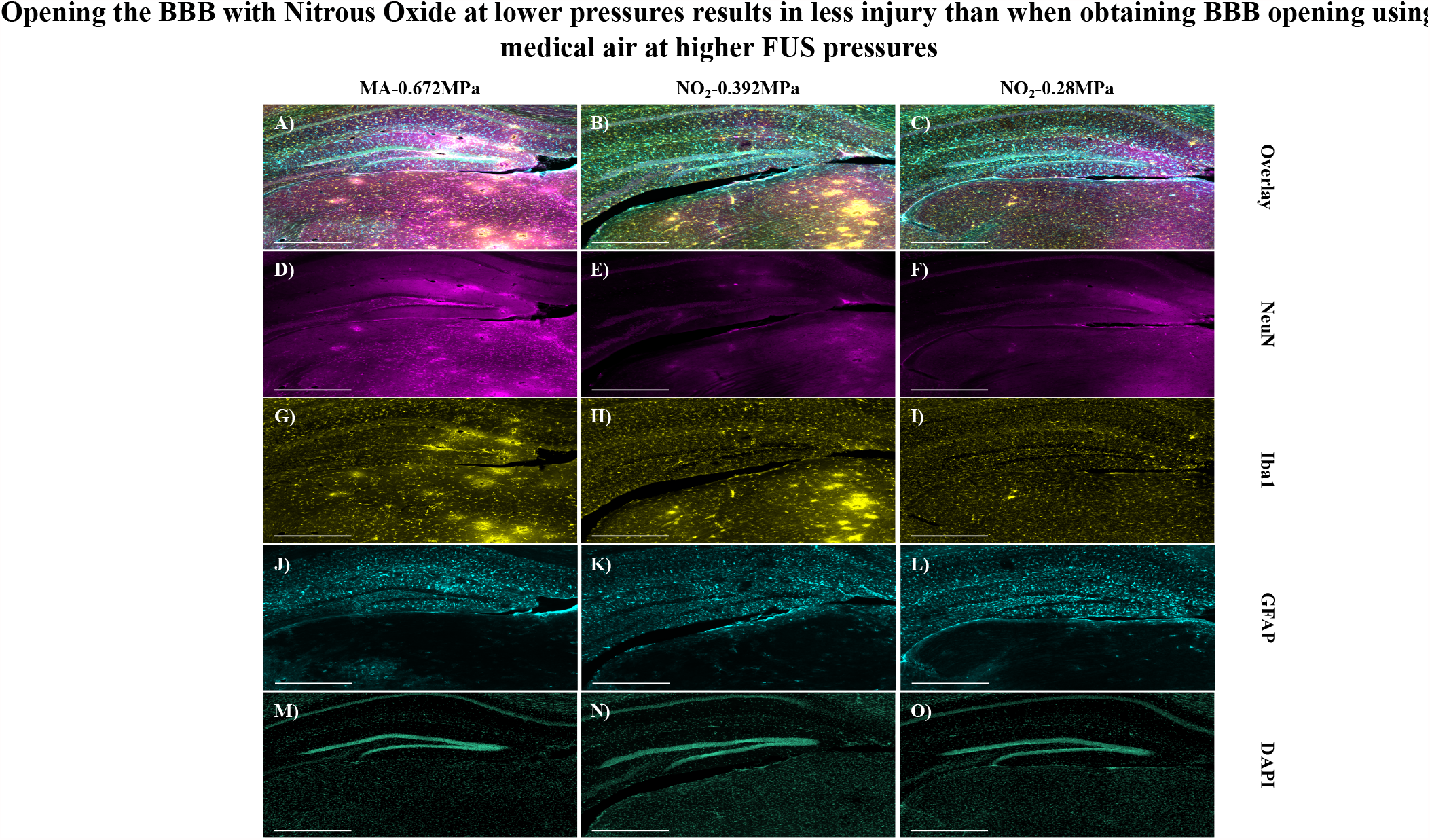
Opening the BBB with Nitrous Oxide at lower pressures results in less injury than when obtaining BBB opening using medical air at higher FUS pressures. A-O) Immunofluorescent stains of coronal brain sections with NeuN (magenta), Iba1 (yellow), GFAP (cyan), and DAPI (blue-green) following FUS mediated BBB opening.A,D,G,J,M) FUS treatment at 0.672 MPa was facilitated by Medical Air (MA). B,E,G,K,N) FUS treatment at 0.392 MPa was facilitated by Nitrous Oxide (NO_2_). C,F,I,L,O) FUS treatment at 0.28 MPa was facilitated by Nitrous Oxide (NO_2_). D-F) The non-specific binding of NeuN indicates extravascular IgG and suggests BBB opening in each brain. G-I) The lack of microgliosis in the NO_2_-0.28MPa treatment group indicates BBB opening was not accompanied by damage. A 500µm scale bar is present in bottom left of each panel.

## Discussion

We have observed that inhaled N_2_O amplifies the acoustic emissions from stimulated intravascular microbubbles required for safe BBB opening and thereby lowers the FUS pressures required. By administering N_2_O, the microbubble dose required to achieve BBB opening can be lowered to at least 1/100 of the current clinical dose. N_2_O potentiates BBB opening in a clinically relevant manner as it substantially improves the delivery of AAV9 to discrete, targeted regions. These findings expand the use of transcranial FUS to non-central brain regions by modulating FUS pressures and microbubble doses for specific targets.

Previous studies from our group and others have demonstrated the ability to deliver a variety of diagnostic and therapeutic agents across the BBB in a targeted manner using FUS and intravenous microbubbles.(*3, 5, 6, 27, 28*) The interaction between the microbubble agent and the ultrasound pressure field governs the efficiency of BBB opening. Microbubble infusions achieve more consistent BBB opening than bolus injections.(*27*) However, if multiple targets are desirable, treatments can be limited by microbubble dose, FUS pressures, and time constraints. Additionally, depletion of the intravascular bubble concentration reduces the signal to noise from acoustic emissions, hindering real-time measurements of stimulated microbubble activity in the brain. Studies examining the utility of monitoring real-time microbubble acoustic emissions have focused on adjusting ultrasound exposure parameters including acoustic pressure, pulse length, and repetition frequencies(*14, 29, 30*) or microbubble parameters such as bubble size or formulation(*6, 25, 31, 32*) to modulate BBB disruption. This assumes that interaction between microbubbles and the brain microvasculature triggers BBB opening, and therefore stabilizing bubble oscillation will trigger opening without damage. This idea is supported by a growing number of preclinical and clinical studies(*29, 31-33*).

Understanding the complete mechanism by which N_2_O potentiates FUS mediated BBB opening will need further study. However, the mechanism of action by which N_2_O amplifies acoustic emissions from stimulated intravascular microbubbles is likely related to a previously observed phenomenon, in which it causes expansion of enclosed gas bubbles. This is a result of 2 factors: i) N_2_O has low potency which results in high blood concentrations and ii) N_2_O has a low blood-gas partition coefficient. The combination of these factors results in N_2_O rapidly moving into confined air spaces (such as a microbubble). The rate at which N_2_O enters a microbubble is faster than the rate at which air moves out, this results in microbubble expansion even before FUS is applied. (*34*) This phenomenon has previously contributed to complications in patients who underwent intraocular gas placement during vitreoretinal surgery and subsequently underwent anesthesia with N_2_O before the intraocular gas bubble had resorbed.(*35*)This mechanism of action is also supported by an early observation in our experiments. When we administered N_2_O without microbubbles we were able to detect subharmonic acoustic emissions in the absence of FUS. However, we did not see BBB opening in the absence of microbubbles. These findings suggest that microbubbles are required for safe BBB opening and that N_2_O merely potentiates the oscillation of the microbubbles thereby reducing the FUS pressure to open the BBB. In addition to the benefits of N_2_O for therapeutic ultrasound application, the impact of this method of bubble response amplification deserves study for diagnostic ultrasound. The ability to improve the harmonic response of microbubbles could lead to improving contrast-enhanced ultrasound imaging. To date, we are unaware of any investigations that have explored this combination of technology.

Global vascular dynamics may also play a large role in the acoustic response observed. This has been suggested in previous work comparing the effect on the acoustic emission of oxygen versus medical air as the delivery gas for anesthesia during in vivo experiments(*18, 31*). Medical air was shown to have a greater occurrence and strength of broadband acoustic emission at higher pressures when compared to oxygen(*31*). Pure oxygen increases the partial pressure of oxygen within the blood resulting in a greater ventilation/perfusion mismatch and increased driving force for microbubble diffusion(*18*) and theoretically a decrease in cerebral blood flow from oxygen related vasoconstriction(*16*). However, N_2_O has the opposite effect and is a known potent cerebral vessel vasodilator. This enables the delivery of more microbubbles to the targeted region, allowing us to decrease the microbubble dose while improving BBB opening.

Previous studies by our group and others show that the use of FUS to deliver intravenously injected AAV9 to targeted regions of the brain allows for doses up to 100 times lower than those conventionally used. Using a 100-fold lower AAV dose, we found that the FUS-mediated potentiation of AAV targeting to the brain with medical air is pressure dependent where higher FUS pressures resulted in a greater number of transduced neurons. However, we also found that higher FUS pressures are associated with increased gliosis, cautioning against the use of higher FUS pressures. To overcome this concern, we demonstrate that N_2_O can be used in combination with a low FUS pressure to effectively target AAV9 vector to the hippocampus, resulting in a greater number of transduced neurons as compared to use of medical air at higher FUS pressures. As such with the use of N_2_O during FUS, it may be feasible to decrease AAV doses lower than previously reported and still achieve high transgene expression. Lower viral doses are of great importance for human translation of gene therapies as higher viral doses result in greater immune responses. Additionally, lower viral doses to achieve an equivalent level of therapeutic benefit could alleviate the high costs and long wait times currently associated with manufacturing human grade AAV vectors.

Transcranial MRgFUS represents a unique translational opportunity to non-invasively validate our understanding of neural networks and to diagnose and treat complex diseases of the brain at the molecular level. The ability to disrupt the BBB in a targeted manner is essential to preventing off-site adverse effects, reducing the cost of therapeutics, and improving efficacy. Although early clinical trials have shown that it is safe to temporarily open the BBB in humans, optimization of parameters to improve the interactions of microbubbles with focused ultrasound will be a critical step in successful human translation. This paper represents an important milestone in that direction by demonstrating the added value of using N_2_O during FUS mediated BBB opening.

### Steps for clinical translation

N_2_O is already a commonly used outpatient analgesic. Early safety and feasibility studies have already shown that FUS mediated opening the BBB in humans is safe, repeatable, and reproducible. As such, this work could very rapidly be translated into the clinic. It would allow for substantial decreases in microbubble dose concentrations and FUS energies while enabling reproducible BBB opening. This would rapidly improve delivery of therapeutics. It would also have the desirable effect of analgesia in patients undergoing focused ultrasound.

### Limitations

There are some limitations to this study. First, the concentration of N_2_O that leaves the typical anesthesia system is not identical to that inspired or that reaching the alveoli. Leakage of N_2_O from components of the anesthesia system is a key factor in that discrepancy.(*5*) As such it is unclear what the actual concentration delivered to alveoli is. Despite this limitation, N_2_O is commonly used in outpatient clinical settings. Second, this study was only performed in mice and should be replicated in larger animal models prior to humans. Finally, the ultrasound frequency used for opening the BBB in this study was 1.5 MHz as the focal dimensions were optimized for a small mouse brain. Although we expect the results to be similar at 0.25 MHz it should be confirmed.

## Materials and Methods

### Study Design

This was a controlled laboratory experiment whose primary pre-specified research objectives were to determine if nitrous oxide could reduce the amount of FUS pressure required to open the BBB in mice and whether a lower concentration of microbubbles could be used. A hypothesis that developed after initiation of data analyses was if the BBB could be opened without microbubbles in the presence of nitrous oxide.

### Animal Studies

Animal protocol 101517 was reviewed and approved for this study by the internal animal ethics committee, University of Texas Southwestern Medical Center Institutional Animal Care and Use Committee (IACUC). Swiss Webster mice (Charles River, Wilmington, MA, USA. n=26) of both genders ages 3 to 7 months were utilized. For viral delivery experiments C57BL/6J mice (males and females, 8-10 weeks old, average weight of 20 g, n=6) were used.

### Experimental design

Twenty-six Swiss Webster mice were randomly selected to undergo an optimization study for the gas and bubble concentration. Twenty were sonicated using Nitrous oxide (N_2_O) and six using Medical air (MA). For the N_2_O cohort, four subcohorts existed. One, using clinical microbubble dose concentration with different pressure 0.39, 0.45 and 0.67 MPa (n=12, 6 with N_2_O and 6 with MA). Second, using clinical microbubble dose concentration with very low pressure 0.28 MPa (n=2). Third, using different bubble concentrations from clinical dose to 1000X dilution at 0.39 MPa (n=6). Fourth, using 1000X dilution of Definity with different pressure from 0.39 to 0.67 MPa (n=6). For viral delivery experiments C57BL/6J mice were selected and three sets of experiments were performed. First with N2O and 0.39 MPa (n=2), Second with MA and 0.39 MPa (n=2), and third with MA and 0.67 MPa (n=2).

### Materials

#### Microbubbles (MB)

Definity (Perflutren Lipid Microsphere) ultrasound contrast agent was purchased from Lantheus Medical Imaging (N. Billerica, MA, USA).

#### Gas Agents

Nitrous Oxide (Airgas, Radnor, PA, USA), Oxygen (Airgas, Radnor, PA, USA). The ratio of N_2_O and O_2_ was preset at 1:3 and the flow rate was 3 LPM using a clinical nitrous oxide delivery system (Belmed, PC-7 flowmeter Nitrous Oxide/Oxygen Sedation, Red Lion, PA, USA). Anesthesia was induced and maintained using 1-3% isoflurane. During animal preparation and MR imaging, anesthesia was maintained with isoflurane mixed into room air supplied by a compressor at 50 PSI and 1 LPM (PM15, Precision Medical, Northampton PA). During focused ultrasound exposures, isoflurane was mixed with the nitrous oxide and oxygen mixture.

#### Air compressor

PM15 EasyAir Compressor (Precision Medical, Inc. Northampton, PA, USA)

### Focused Ultrasound System

The ultrasound system used to deliver FUS exposures to the mouse brain was a stereotactic atlas-guided system (RK-50, FUS Instruments, Toronto, ON, CA). A custom-built Python-based software suite enabled landmark localization, creation of a treatment plan on a representation of a mouse brain, and execution of the treatment through control of the systems motors and ultrasonic field. The accompanying focused ultrasound transducer operated at 1.43 MHz and had a 35 mm diameter and 24.5 mm radius of curvature. This frequency was utilized to obtain spatial localization of ultrasound within the mouse brain. The pressure output and beam profile of the transducer was calibrated with a needle hydrophone (Precision Acoustics, Dorchester, UK).

### Animal Preparation

Swiss Webster mice (Charles River, Wilmington, MA, USA. n=26) of both genders ages 3 to 7 months were utilized. Each animal was anesthetized initially through inhalation using a mixture of 1.5-3% isoflurane and 1 L/min of Medical Air. SuperCath 5, 26G Safety IV Catheter (ICU Medical, Inc. CA, USA) closed with injection cap with a measured dead volume of 130 µL was placed in the lateral tail vein for bubble and therapy administration. A physiologic monitoring system (PhysioSuite, Kent Scientific Corp., Torrington, CT, USA) was used to monitor vital signs and maintain core body temperature throughout the experiment. Hair over the cranial surface of the skull was removed using an animal clipper and depilatory cream (VEET sensitive formula, Reckitt Benckiser, Parsippany, NJ, USA). Using sterile technique, a 1-2 cm incision was performed over the skull to visualize bregma and lambda skull suture landmarks. The animal’s head was stabilized on the stereotaxic apparatus using ear bars and a bite bar. A nose cone was placed over the animal’s nose to deliver inhalant anesthetic during sonication. Once positioned on the stereotactic unit, the inhalant gas of interest was administered for 15 min at 1 L/min of Medical Air or 3 L/min 70% Nitrous Oxide mixed with 30% Oxygen with 1.5-2% isoflurane prior to starting sonication. Once the skull landmarks were established, ultrasound gel was added to the animal’s skull for acoustic coupling. A tank filled with deionized and degassed water was lowered onto the ultrasound gel, and the transducer was lowered. For viral delivery experiments C57BL/6J mice (males and females, 8-10 weeks old, average weight of 20 g, n=2 per treatment group) were used, and were prepared similarly to the swiss-webster mice. All experiments were performed with the approval of the Institutional Animal Care and Use Committee of the University of Texas Southwestern Medical Center. All methods and protocols were carried out in accordance with the guidelines set forth in the Guide for the Care and Use of Laboratory Animals.

### Virus Preparation

A reporter vector was used in these studies where gene expression cassette encoded eGFP under the control of the CBh promoter (REF) and with a SV40pA. Self-complementary AAV vector was produced using methods developed by the University of North Carolina (UNC) Vector Core facility, as described. The purified AAV was dialyzed in PBS supplemented with 5% D-Sorbitol and an additional 212 mM NaCl (350 mM NaCl total). Vector was titered by quantitative PCR,(*36*) and confirmed by polyacrylamide gel electrophoresis and silver stain. AAV9 vector was diluted in the vehicle and each animal received a dose of 1×10^9^ vg/g by tail vein infusion following the FUS-BBB opening procedure.

### Bubble Preparation and Infusion

Definity MB were activated using the shaking amalgamator provided by the manufacturer for 45 sec for IV infusion, manufacturer recommendations for human use specify a dilution of 1.5 µL of Definity added to 50 mL of preservative-free saline delivered at a rate between 4 mL/minute to 10 mL/minute. This dosing differs significantly from the instructions for bolus administration, which is recommended at a dose of 10 µL per kg over 30 to 60 seconds. For an average human weight of 80 kg, the Definity delivered will be approximately 800 µL per a single bolus. However, over the length of an infusion, the Definity delivered will be greater. Since the majority of pre-clinical studies performed utilize bolus delivery and a dosage range of 10 to 20 µL per kg, we chose to use that as a basis for our infusion dose by calculating a dose that delivers 20 µL per kg animal weight per sonication location over the course of an infusion. 3.88 ul of microbubbles were drawn from the prepared microbubble vial, and were diluted in 596.12 ul of saline. The diluted microbubble mixture was infused into the animal at a rate of 50 ul/min using an infusion pump (Nanojet, Chemyx Inc, Stafford, TX, USA), through the intravenous catheter inserted into a tail vein. The infusion ran during the focused ultrasound exposure. FUS exposures were commenced after 30 µL of the bubble solution had entered circulation.

### Sonication Treatment Plans

#### AUC and Postcontrast MRI Studies

The sonication targets were: Striatum, Hippocampus and Cerebellum. At each target 1-6 exposures were performed. Each exposure consisted of 20 bursts with a pulse length of 10 ms and a repetition period of 1000 ms. The pressure applied at target locations was 0.392, 0.448 & 0.672 (MPa).

#### Viral Gene Therapy Delivery Studies

The sonication targets were Striatum, Hippocampus and Cerebellum at each target 1-2 exposures were performed separated by 2mm. Each exposure consisted of 20 bursts with a pulse length of 10 ms and a repetition period of 1000 ms. The pressures applied at target locations in the right hemisphere were 0.392 for Nitrous Oxide and 0.672 (MPa) for medical air.

##### Data Collection

Acoustic Emissions: The acoustic emissions from microbubbles stimulated at the ultrasound focus were captured by a hydrophone integrated into the center of the transducer and digitized by the FUS system using an on-board oscilloscope (Picoscope 5242D, Pico Technologies, United Kingdom). The hydrophone had a resonant frequency of approximately 700 kHz and was selected to match the subharmonic frequency of microbubbles stimulated at 1.43 MHz. The received hydrophone signal after each burst was processed to calculate the frequency spectrum of this signal in MATLAB. A unique frequency spectrum was calculated for each burst, resulting in 20-30 spectra for each brain location. To compare the frequency response across different experiment conditions, an area under the curve (AUC) was defined as the sum of the harmonic amplitudes in the frequency spectra centered at the subharmonic of the transmit frequency for all bursts at a single pressure and in a single exposure location. The frequency band from 714 – 716 kHz was used to calculate the AUC. Dextran Slides: Images were collected using a Zeiss Axioscan.Z1 (Carl Zeiss AG, Oberkochen, Germany) with the Zen Blue Software suite (Carl Zeiss AG, Oberkochen, Germany)

##### Immunohistochemistry and quantification

Four weeks-post injection, mice were deeply anesthetized with isoflurane and perfused with PBS containing 1 µg/mL heparin and then perfused with 10% formalin. Brains were drop-fixed in 10% formalin for 48 hours before being moved to PBS. Brains were cut down the midline and each half vibratome sectioned into 40 µm sagittal slices using a Leica Vibratome (VT1000S; Leica Biosystems Inc., Buffalo Grove, IL) and stored in PBS at 4°C. Every 5^th^ section from the whole brain of each animal was stained free-floating as previously described (*37*). Primary antibody was anti-GFP (1:1500) (Millipore, 3080) and secondary was anti-rabbit (1:1000) (Vector Laboratories, Burlingame, CA). Images were captured using a Zeiss Axioscan.Z1 with the ImageScope Software (Leica Biosystems Inc., Buffalo Grove, IL).

#### Immunofluorescence

##### Isolation of mouse brain

Animals were anesthetized with isoflurane and perfused with cold PBS. Whole-brains were drop-fixed in Phosphate-Buffered 4% Paraformaldehyde (FD NeuroTechnologies, Colombia, MD, USA) overnight at 4 °C. Brains were then placed in 10% sucrose in PBS for 24 hrs at 4 °C, followed by 24 hrs in 20% sucrose in PBS at 4 °C, and finally stored in 30% sucrose in PBS at 4 °C until sectioning.

##### Sectioning

A sliding-base freezing microtome (Thermo-Fisher Scientific, Waltham, MA, USA) was used to collect 40 µm free-floating coronal sections from fixed mouse brains. Sections were stored in cryoprotectant at 4 °C until IHC performed.

##### Staining

For triple-stain fluorescent-IHC, sections were incubated in Rabbit anti-Iba (1:1000 Wako Chemicals USA, Richmond, VA, USA), Goat anti-GFAP (1:500, abcam, Cambridge, MA, USA), and mouse anti-NeuN (1:200, Millipore, Burlington, MA, USA) overnight at 4 °C. Sections were then washed and incubated with AlexaFluor®488 Donkey anti-rabbit IgG (1 : 500), AlexaFluor®647 Donkey anti-goat IgG (1 : 500), and AlexaFluor®568 Donkey anti-mouse IgG(1 : 500, all from Thermo-Fisher Scientific, Waltham, MA, USA) for 1 hour, followed by 10 minutes in a 1:1000 DAPI solution (Sigma, St.Louis, MO, USA). Sections were rinsed, mounted, allowed to dry overnight, and then cover slipped with Aqua-Poly/Mount (Polysciences Inc., Warrington, PA, USA). Sections were then imaged on a Zeiss Axioscan.Z1 (Zeiss, White Plains, NY, USA) in the Univ of Texas Southwestern Medical Center Whole Brain Microscopy Core Facility, RRID:SCR_017949). Slides were imaged at 20X using the same settings for exposure time and intensity between slides. Control sections from mice not receiving treatment were utilized to confirm staining with and without primary antibodies.

#### Data Analysis

##### AUC

To compare the frequency response across different experiment conditions, an area under the curve (AUC) was defined as the sum of the harmonic amplitudes in the frequency spectra centered at the subharmonic of the transmit frequency for all bursts at a single pressure and in a single exposure location. The frequency band from 714 – 716 kHz was used to calculate the AUC.

##### GFP positive neurons

The number of positive hippocampal cells were determined by blinded counts of 3-4 representative hippocampal sections located between 1.96 and 2.5 mm lateral to the midline from both the left and right hemisphere for each animal. The total number of neurons were averaged per number of sections counted per side and per animal. Positive neurons required the presence of a positive soma and were distinguished from positive glia based on morphology.

Statistical analysis of AUCs between experimental conditions were performed using Student t-test where applicable. Statistical analysis of GFP positive neurons were performed using Two-way ANOVA and post hoc Sidak’s multiple comparisons test with treatment condition and brain hemisphere (open versus closed) as independent variables. Analysis was performed using GraphPad Prism version 9.1.0 software (GraphPad Software) and P<0.05 was considered to be statistically significant.

### Ethics Statement

All experiments were performed after approval of the Institutional Animal Care and Use Committee (IACUC) of University of Texas Southwestern and conformed to the National Institutes of Health’s PHS Policy on the Humane Care and Use of Laboratory Animals.

## Funding

UTSW High Impact Award: BRS, RMB, RC

## Author contributions

Conceptualization: BRS, RMB, RC

Methodology: BRS, RMB, RC

Validation: BRS, RMB, SKH, MD, RC

Formal Analysis: BRS, RMB, IY, SKH, MD, RC

Investigation: BRS, RMB, IY, RC

Resources: BRS, RMB, MD, RC

Visualization: BRS, RMB, IY, SKH, AM, SJET

Funding acquisition: BRS, RMB, RC

Project administration: BRS, RMB, RC

Supervision: BRS, RMB, RC

Writing – original draft: BRS, RMB, RC

Writing – review & editing: BRS, RMB, IY, AM, MD, RC

## Competing Interests

RC has a competing interest with FUS instruments. Other authors declare that they have no competing interests.

## Data and materials availability

All data are available in the main text.

## Notes

### Competing Interest Statement

RC - FUS instruments

## References and Notes

1. J. Fenstermacher, J. Gazendam, Intra-arterial infusions of drugs and hyperosmotic solutions as ways of enhancing CNS chemotherapy. Cancer Treat Rep 65 Suppl 2, 27–37 (1981).

2. S. I. Rapoport, M. Hori, I. Klatzo, Testing of a hypothesis for osmotic opening of the blood-brain barrier. Am J Physiol 223, 323–331 (1972).

3. D. S. Hersh, A. S. Wadajkar, N. Roberts, J. G. Perez, N. P. Connolly, V. Frenkel, J. A. Winkles, G. F. Woodworth, A. J. Kim, Evolving Drug Delivery Strategies to Overcome the Blood Brain Barrier. Curr Pharm Des 22, 1177–1193 (2016).

4. W. M. Pardridge, Biopharmaceutical drug targeting to the brain. J Drug Target 18, 157–167 (2010).

5. C. Bing, M. Ladouceur-Wodzak, C. R. Wanner, J. M. Shelton, J. A. Richardson, R. Chopra, Transcranial opening of the blood-brain barrier in targeted regions using a stereotaxic brain atlas and focused ultrasound energy. J Ther Ultrasound 2, 13 (2014).

6. B. Cheng, C. Bing, Y. Xi, B. Shah, A. A. Exner, R. Chopra, Influence of Nanobubble Concentration on Blood-Brain Barrier Opening Using Focused Ultrasound Under Real-Time Acoustic Feedback Control. Ultrasound Med Biol 45, 2174–2187 (2019).

7. N. McDannold, N. Vykhodtseva, S. Raymond, F. A. Jolesz, K. Hynynen, MRI-guided targeted blood-brain barrier disruption with focused ultrasound: histological findings in rabbits. Ultrasound Med Biol 31, 1527–1537 (2005).

8. D. E. Becker, M. Rosenberg, Nitrous oxide and the inhalation anesthetics. Anesth Prog 55, 124–130; quiz 131-122 (2008).

9. T. Kobus, N. Vykhodtseva, M. Pilatou, Y. Zhang, N. McDannold, Safety Validation of Repeated Blood-Brain Barrier Disruption Using Focused Ultrasound. Ultrasound Med Biol 42, 481–492 (2016).

10. G. Samiotaki, F. Vlachos, Y. S. Tung, E. E. Konofagou, A quantitative pressure and microbubble-size dependence study of focused ultrasound-induced blood-brain barrier opening reversibility in vivo using MRI. Magn Reson Med 67, 769–777 (2012).

11. M. A. O’Reilly, K. Hynynen, Blood-brain barrier: real-time feedback-controlled focused ultrasound disruption by using an acoustic emissions-based controller. Radiology 263, 96–106 (2012).

12. C. H. Tsai, J. W. Zhang, Y. Y. Liao, H. L. Liu, Real-time monitoring of focused ultrasound blood-brain barrier opening via subharmonic acoustic emission detection: implementation of confocal dual-frequency piezoelectric transducers. Phys Med Biol 61, 2926–2946 (2016).

13. Y. S. Tung, F. Marquet, T. Teichert, V. Ferrera, E. E. Konofagou, Feasibility of noninvasive cavitation-guided blood-brain barrier opening using focused ultrasound and microbubbles in nonhuman primates. Appl Phys Lett 98, 163704 (2011).

14. C. D. Arvanitis, M. S. Livingstone, N. Vykhodtseva, N. McDannold, Controlled ultrasound-induced blood-brain barrier disruption using passive acoustic emissions monitoring. PLoS One 7, e45783 (2012).

15. K. B. Bader, C. K. Holland, Gauging the likelihood of stable cavitation from ultrasound contrast agents. Phys Med Biol 58, 127–144 (2013).

16. N. McDannold, Y. Zhang, N. Vykhodtseva, Blood-brain barrier disruption and vascular damage induced by ultrasound bursts combined with microbubbles can be influenced by choice of anesthesia protocol. Ultrasound Med Biol 37, 1259–1270 (2011).

17. N. McDannold, Y. Zhang, J. G. Supko, C. Power, T. Sun, C. Peng, N. Vykhodtseva, A. J. Golby, D. A. Reardon, Acoustic feedback enables safe and reliable carboplatin delivery across the blood-brain barrier with a clinical focused ultrasound system and improves survival in a rat glioma model. Theranostics 9, 6284–6299 (2019).

18. E. Sassaroli, K. Hynynen, Resonance frequency of microbubbles in small blood vessels: a numerical study. Phys Med Biol 50, 5293–5305 (2005).

19. A. Qamar, R. Samtaney, J. L. Bull, Dynamics of micro-bubble sonication inside a phantom vessel. Appl Phys Lett 102, 13702 (2013).

20. C. Chen, Y. Gu, J. Tu, X. Guo, D. Zhang, Microbubble oscillating in a microvessel filled with viscous fluid: A finite element modeling study. Ultrasonics 66, 54–64 (2016).

21. M. J. Cipolla, in The Cerebral Circulation. (San Rafael (CA), 2009).

22. N. McDannold, Y. Zhang, N. Vykhodtseva, The Effects of Oxygen on Ultrasound-Induced Blood-Brain Barrier Disruption in Mice. Ultrasound Med Biol 43, 469–475 (2017).

23. L. Mullin, R. Gessner, J. Kwan, M. Kaya, M. A. Borden, P. A. Dayton, Effect of anesthesia carrier gas on in vivo circulation times of ultrasound microbubble contrast agents in rats. Contrast Media Mol Imaging 6, 126–131 (2011).

24. N. Hosseinkhah, K. Hynynen, A three-dimensional model of an ultrasound contrast agent gas bubble and its mechanical effects on microvessels. Phys Med Biol 57, 785–808 (2012).

25. J. J. Choi, K. Selert, Z. Gao, G. Samiotaki, B. Baseri, E. E. Konofagou, Noninvasive and localized blood-brain barrier disruption using focused ultrasound can be achieved at short pulse lengths and low pulse repetition frequencies. J Cereb Blood Flow Metab 31, 725–737 (2011).

26. G. Samiotaki, E. E. Konofagou, Dependence of the reversibility of focused-ultrasound-induced blood-brain barrier opening on pressure and pulse length in vivo. IEEE Trans Ultrason Ferroelectr Freq Control 60, 2257–2265 (2013).

27. C. Bing, Y. Hong, C. Hernandez, M. Rich, B. Cheng, I. Munaweera, D. Szczepanski, Y. Xi, M. Bolding, A. Exner, R. Chopra, Characterization of different bubble formulations for blood-brain barrier opening using a focused ultrasound system with acoustic feedback control. Sci Rep 8, 7986 (2018).

28. Y. S. Tung, F. Vlachos, J. J. Choi, T. Deffieux, K. Selert, E. E. Konofagou, In vivo transcranial cavitation threshold detection during ultrasound-induced blood-brain barrier opening in mice. Phys Med Biol 55, 6141–6155 (2010).

29. A. Abrahao, Y. Meng, M. Llinas, Y. Huang, C. Hamani, T. Mainprize, I. Aubert, C. Heyn, S. E. Black, K. Hynynen, N. Lipsman, L. Zinman, First-in-human trial of blood-brain barrier opening in amyotrophic lateral sclerosis using MR-guided focused ultrasound. Nat Commun 10, 4373 (2019).

30. Y. S. Tung, F. Vlachos, J. A. Feshitan, M. A. Borden, E. E. Konofagou, The mechanism of interaction between focused ultrasound and microbubbles in blood-brain barrier opening in mice. J Acoust Soc Am 130, 3059–3067 (2011).

31. Y. Meng, B. J. MacIntosh, Z. Shirzadi, A. Kiss, A. Bethune, C. Heyn, K. Mithani, C. Hamani, S. E. Black, K. Hynynen, N. Lipsman, Resting state functional connectivity changes after MR-guided focused ultrasound mediated blood-brain barrier opening in patients with Alzheimer’s disease. Neuroimage 200, 275–280 (2019).

32. S. H. Park, M. J. Kim, H. H. Jung, W. S. Chang, H. S. Choi, I. Rachmilevitch, E. Zadicario, J. W. Chang, Safety and feasibility of multiple blood-brain barrier disruptions for the treatment of glioblastoma in patients undergoing standard adjuvant chemotherapy. J Neurosurg, 1–9 (2020).

33. E. S. Lein, M. J. Hawrylycz, N. Ao, M. Ayres, A. Bensinger, A. Bernard, A. F. Boe, M. S. Boguski, K. S. Brockway, E. J. Byrnes, L. Chen, L. Chen, T. M. Chen, M. C. Chin, J. Chong, B. E. Crook, A. Czaplinska, C. N. Dang, S. Datta, N. R. Dee, A. L. Desaki, T. Desta, E. Diep, T. A. Dolbeare, M. J. Donelan, H. W. Dong, J. G. Dougherty, B. J. Duncan, A. J. Ebbert, G. Eichele, L. K. Estin, C. Faber, B. A. Facer, R. Fields, S. R. Fischer, T. P. Fliss, C. Frensley, S. N. Gates, K. J. Glattfelder, K. R. Halverson, M. R. Hart, J. G. Hohmann, M. P. Howell, D. P. Jeung, R. A. Johnson, P. T. Karr, R. Kawal, J. M. Kidney, R. H. Knapik, C. L. Kuan, J. H. Lake, A. R. Laramee, K. D. Larsen, C. Lau, T. A. Lemon, A. J. Liang, Y. Liu, L. T. Luong, J. Michaels, J. J. Morgan, R. J. Morgan, M. T. Mortrud, N. F. Mosqueda, L. L. Ng, R. Ng, G. J. Orta, C. C. Overly, T. H. Pak, S. E. Parry, S. D. Pathak, O. C. Pearson, R. B. Puchalski, Z. L. Riley, H. R. Rockett, S. A. Rowland, J. J. Royall, M. J. Ruiz, N. R. Sarno, K. Schaffnit, N. V. Shapovalova, T. Sivisay, C. R. Slaughterbeck, S. C. Smith, K. A. Smith, B. I. Smith, A. J. Sodt, N. N. Stewart, K. R. Stumpf, S. M. Sunkin, M. Sutram, A. Tam, C. D. Teemer, C. Thaller, C. L. Thompson, L. R. Varnam, A. Visel, R. M. Whitlock, P. E. Wohnoutka, C. K. Wolkey, V. Y. Wong, M. Wood, M. B. Yaylaoglu, R. C. Young, B. L. Youngstrom, X. F. Yuan, B. Zhang, T. A. Zwingman, A. R. Jones, Genome-wide atlas of gene expression in the adult mouse brain. Nature 445, 168–176 (2007).

34. R. Benavides, M. Maze, N. P. Franks, Expansion of gas bubbles by nitrous oxide and xenon. Anesthesiology 104, 299–302 (2006).

35. B. J. Vote, R. H. Hart, D. R. Worsley, J. H. Borthwick, S. Laurent, A. J. McGeorge, Visual loss after use of nitrous oxide gas with general anesthetic in patients with intraocular gas still persistent up to 30 days after vitrectomy. Anesthesiology 97, 1305–1308 (2002).

36. S. J. Gray, V. W. Choi, A. Asokan, R. A. Haberman, T. J. McCown, R. J. Samulski, Production of recombinant adeno-associated viral vectors and use in in vitro and in vivo administration. Curr Protoc Neurosci Chapter 4, Unit 4 17 (2011).

37. R. M. Bailey, A. Rozenberg, S. J. Gray, Comparison of high-dose intracisterna magna and lumbar puncture intrathecal delivery of AAV9 in mice to treat neuropathies. Brain Res 1739, 146832 (2020).

